## Supplementary Materials for "*Cdkn2a* regulates beige fat maintenance through BECN1-mediated autophagy"

**through BECN1-mediated autophagy**

**Figure Legend**

**Figure S1. Knockout of *Cdkn2a* does not affect BAT characteristics after the rewarming period following cold exposure**

(A) NMR analysis of fat (left panel) and lean masses (right panel) of body weight of control or *Cdkn2a^Ucp1^* KO mice fed a chow diet.
(B) Representative H&E staining images of BAT from control or *Cdkn2a^Ucp1^* KO mice. Scale bar, 100 μM.
(C) Immunofluorescence staining of UCP1 and RFP in BAT from control or *Cdkn2a^Ucp1^* KO mice. Scale bar, 100 μM.
(D) Quantification of the percentage of RFP^+^ cells that express endogenous UCP1.
(E) Western blot analysis of UCP1 in BAT from control or *Cdkn2a^Ucp1^* KO mice.
(F) qPCR analysis of mRNA expression of thermogenic genes in BAT from control or *Cdkn2a^Ucp1^* KO mice at day 0 (left panel) and day 60 (right panel) post withdrawal cold stimulus.
(G) Representative H&E staining images of liver from control or *Cdkn2a^Ucp1^* KO mice. Scale bar, 100 μM.
(H) Tile scan of IGW and PGW from control or *Cdkn2a^Ucp1^* KO mice. Scale bar, 100 μM.
(I) Quantification of adipocyte sizes in PGW from control or *Cdkn2a^Ucp1^* KO mice.
(J) Weight of adipose tissues and other tissues from control or *Cdkn2a^Ucp1^* KO mice.

^*^p < 0.05, ^**^p < 0.01, ^***^p < 0.001 by two-tailed Student’s t test. n = 6-8 mice per group. Data are expressed as means ± SEM.

**Figure S2. *Cdkn2a* deficiency has no effect on BAT thermogenesis in mice under HFHS**

(A) Food intake of control or *Cdkn2a^Ucp1^* KO mice under HFHS.
(B) Weight of adipose tissues and other tissues from control or *Cdkn2a^Ucp1^* KO mice under HFHS.
(C) Tile scan of PGW from control or *Cdkn2a^Ucp1^* KO mice under HFHS. Scale bar, 100 μM.
(D) Quantification of adipocyte sizes in PGW from control or *Cdkn2a^Ucp1^* KO mice.
(E) Representative H&E staining images of BAT and liver from control or *Cdkn2a^Ucp1^* KO mice under HFHS. Scale bar, 100 μM.
(F) O_2_ consumption of control or *Cdkn2a^Ucp1^* KO mice fed a HFHS. White and gray areas in the graphs indicate light and night, respectively.
(G) CO_2_ generation of control or *Cdkn2a^Ucp1^* KO mice fed a HFHS.
(H) Locomotor activity of control or *Cdkn2a^Ucp1^* KO mice fed a HFHS.
(I) Immunofluorescence staining of UCP1 and RFP in BAT from control or *Cdkn2a^Ucp1^* KO mice under HFHS. Scale bar, 50 μM.
(J) Quantification of the percentage of RFP^+^ cells that express endogenous UCP1.
(K) Immunohistochemical detection of UCP1 in IGW from control or *Cdkn2a^Ucp1^* KO mice under HFHS. Scale bar, 100 μM.
(L) qPCR analysis of mRNA expression of thermogenic genes in BAT from control or *Cdkn2a^Ucp1^* KO mice under HFHS.

^*^p < 0.05, ^**^p < 0.01, ^***^p < 0.001 by two-tailed Student’s t test. n = 6-8 mice per group. Data are expressed as means ± SEM.

**Figure S3. *Cdkn2a* is associated with beige adipocyte maintenance**

(A) qPCR analysis of the mRNA levels of *p16^Ink4a^* and *p19^Arf^* in brown, beige and white adipocytes.
(B) Expression profile of *p16^Ink4a^* and *p19^Arf^* during beige adipocyte maintenance.
(C) qPCR analysis of mRNA expression of *p16^Ink4a^*, *p19^Arf^* and thermogenic genes in control and *Cdkn2a^Ucp1^* KO beige adipocytes before withdrawing external stimuli.

^*^p < 0.05, ^**^p < 0.01, ^***^p < 0.001 by two-tailed Student’s t test. Data are expressed as means ± SEM of triplicate tests.

**Figure S4. *Cdkn2a* knockout does not affect brown fat maintenance *in vitro***

(A) Schematic illustration of the cellular system of brown adipocyte maintenance. SVF cells isolated from BAT of control or *Cdkn2a^Ucp1^* KO mice were differentiated into brown adipocytes and treated with 4-OHT to induce gene deletion and RFP labeling. Then, external stimuli were withdrawn by changing the differentiation medium to DMEM containing 5% BSA for 4 days.
(B) Expression profile of *p16^Ink4a^* and *p19^Arf^* during brown adipocyte maintenance.
(C) Bright field images of control and *Cdkn2a^Ucp1^* KO brown adipocytes during brown adipocyte maintenance. Scale bar, 100 μM.
(D) Immunofluorescence staining of UCP1 and RFP in control and *Cdkn2a^Ucp1^* KO brown adipocytes during brown adipocyte maintenance. Scale bar, 50 μM.
(E) Quantification of the percentage of RFP+ cells that express endogenous UCP1.
(F) qPCR analysis of the mRNA expression of *p16^Ink4a^*, *p19^Arf^*, and thermogenic genes in control and *Cdkn2a^Ucp1^* KO brown adipocytes before withdrawing external stimuli.
(G) qPCR analysis of the mRNA expression of thermogenic genes in control and *Cdkn2a^Ucp1^* KO brown adipocytes during brown adipocyte maintenance.

^*^p < 0.05, ^**^p < 0.01, ^***^p < 0.001 by two-tailed Student’s t test. Data are expressed as means ± SEM of triplicate tests.

**Figure S5. CCND1 does not regulate beige adipocyte maintenance**

(A) Immunofluorescence staining of BrdU and RFP in control and *Cdkn2a^Ucp1^* KO beige adipocytes after withdrawing external stimuli. Scale bar, 100 μM.
(B) Expression profile of *Ccnd1* during beige adipocyte maintenance.
(C) Scheme of *Adipoq* promoter-driven Dox-inducible *Ccnd1* transgenic mice.
(D) Schematic illustration of the cellular system. SVF cells isolated from IGW of control or *Ccnd1^Adipoq^* OE mice were differentiated into beige adipocytes and treated with doxycycline (Dox) to induce gene overexpression. Then, external stimuli were withdrawn by changing the differentiation medium to DMEM containing 5% BSA for 4 days.
(E) qPCR analysis of mRNA levels of *Ccnd1* in control and *Ccnd1^Adipoq^* OE beige adipocytes.
(F) Immunofluorescence staining of UCP1 in control and *Ccnd1^Adipoq^* OE beige adipocytes during beige adipocyte maintenance. Scale bar, 50 μM.
(G) Western blot analysis of UCP1 and LC3 in control and *Ccnd1^Adipoq^* OE beige adipocytes during beigeadipocyte maintenance.

^*^p < 0.05, ^**^p < 0.01, ^***^p < 0.001 by two-tailed Student’s t test. Data are expressed as means ± SEM of triplicate tests.

**Figure S6. Becn1 is a potential regulator of beige fat maintenance**

(A) qPCR analysis of the mRNA expression of autophagy-related genes in control and *Cdkn2a^Ucp1^* KO beige adipocytes after withdrawing external stimuli.
(B) Western blot analysis of BECN1 in control and *Cdkn2a^Ucp1^* KO beige adipocytes after withdrawing external stimuli.
(C) qPCR analysis of the mRNA levels of *Becn1* in IGW or BAT from wild type mice.
(D) qPCR analysis of the mRNA levels of *Becn1* in brown, beige, and white adipocytes.
(E) qPCR analysis of the mRNA expression of *Becn1* in BAT from control or *Cdkn2a^Ucp1^* KO mice (n = 6).
(F) qPCR analysis of *Atg5*, *Atg7* and *Becn1* mRNA expression in control and *Cdkn2a^Ucp1^* KO brown adipocytes after withdrawal of external stimuli.
(G) Scheme of the autophagy-hyperactive *Becn1*^F121A^ knock-in mouse model.

^*^p < 0.05, ^**^p < 0.01, ^***^p < 0.001 by two-tailed Student’s t test. Data are expressed as means ± SEM of triplicate tests.

**Figure S7. Hyperactive *Becn1*^F121A^ does not influence brown adipocyte differentiation and maintenance.**

(A) Schematic illustration of the cellular system of brown adipocyte maintenance. SVF cells isolated from BAT of control or *Becn1*^F121A^ mice were differentiated into brown adipocytes. Then, external stimuli were withdrawn by changing the differentiation medium to DMEM containing 5% BSA for 2 days.
(B) Immunofluorescence staining of LC3 in control and *Becn1*^F121A^ brown adipocytes. Scale bar, 10 μM.
(C) Quantification of the number of LC3 puncta per cell.
(D) Oil red O staining of control and *Becn1*^F121A^ brown adipocytes before withdrawing external stimuli. Scale bar, 100 μM.
(E) Relative lipid accumulation was quantified with a microplate spectrophotometer.
(F) Immunofluorescence staining of UCP1 and PLIN1 in control and *Becn1*^F121A^ brown adipocytes during brown adipocyte maintenance. Scale bar, 50 μM.
(G) Quantification of the percentage of UCP1-expressing cells.
(H) qPCR analysis of mRNA expression of thermogenic genes in control and *Becn1*^F121A^ brown adipocytes after withdrawal of external stimuli.

^*^p < 0.05, ^**^p < 0.01, ^***^p < 0.001 by two-tailed Student’s t test. Data are expressed as means ± SEM of triplicate tests.

**Figure S8. *Cdkn2a* and *Becn1* expression levels positively correlate with obesity in mice and humans**

(A) qPCR analysis of the mRNA expression of *p16^Ink4a^*, *p19^Arf^* and *Becn1* in IGW from mice fed with chow diet or HFHS.
(B) qPCR analysis of the mRNA expression of *p16^Ink4a^*, *p14^Arf^* and *Becn1* in subcutaneous WAT from obese and non-obese people.
(C) Pearson's correlation analysis of *p16^INK4a^* expression in human subcutaneous WAT and body weight, BMI or fat mass of 17 individuals.
(D) Pearson's correlation analysis of *p14^ARF^* expression in human subcutaneous WAT and body weight, BMI or fat mass of 17 individuals.
(E) Pearson's correlation analysis of *BECN1* expression in human subcutaneous WAT and body weight, BMI, fat mass and blood glucose levels of 15-17 individuals.

**Table S1, Related to STAR Methods.**

**Sequences of primers.**

| **Gene** | **Species** | **Forward primer (5’-3’)** | **Reverse primer (5’-3’)** |
| --- | --- | --- | --- |
| *Adipoq* | Mouse | GCAGGCATCCCAGGACATC | GCGATACATATAAGCGGCTTCT |
| *Atg5* | Mouse | ATGCGGTTGAGGCTCACTTTA | GGTTGATGGCCCAAAACTGG |
| *Atg12* | Mouse | TGTGAATCAGTCCTTTGCCCC | TGCAGGACCAGTTTACCATCAC |
| *Becn1* | Mouse | ATGCAGGTGAGCTTCGTGTG | AATGGCTCCTGTGAGTTCCTG |
| *Bnip3* | Mouse | TCCTGGGTAGAACTGCACTTC | GCTGGGCATCCAACAGTATTT |
| *Bnip3l* | Mouse | TGTCTCACTTAGTCGAGCCGC | TGGGTAGCTCCACCCAGGAA |
| *Cdkn2a (p16^Ink4a^)* | Mouse | CGAACTCTTTCGGTCGTAC | ATCATCATCACCTGAATCGGGGT |
| *Cdkn2a (p19^Arf^)* | Mouse | GCAGGTTCTTGGTCACTGT | TCGCACGAACTTCACC |
| *Cidea* | Mouse | TGACATTCATGGGATTGCAGAC | GGCCAGTTGTGATGACTAAGAC |
| *Fabp4* | Mouse | CTGGGCGTGGAATTCGAT | GCTCTTCACCTTCCTGTCGTCT |
| *Fundc1* | Mouse | ACCATAGACTTCCAACTCGGC | TTCCGGGATGCCATGATACC |
| *Leptin* | Mouse | AAGACCATTGTCACCAGGATCAA | GGATACCGACTGCGTGTGTG |
| *Pink1* | Mouse | CCACCTTTCCCTTTGCCATC | ACTGCTCCATACTCTCCAGC |
| *Prdm16* | Mouse | ACACGCCAGTTCTCCAACCTGT | TGCTTGTTGAGGGAGGAGGTA |
| *Pparg* | Mouse | GCATGGTGCCTTCGCTGA | TGGCATCTCTGTGTCAACCATG |
| *Ppargc1a* | Mouse | CCGATCACCATATTCCAGGT | GTGTGCGGTGTCTGTAGTGG |
| *Rrlp0* | Mouse | TCCAGGCTTTGGGCATCA | CTTTATCAGCTGCACATCACTCAGA |
| *Ucp1* | Mouse | CACCTTCCCGCTGGACACT | CCCTAGGACACCTTTATACCTAATGG |
| *ACTB* | Human | GAGCTACGAGCTGCCTGACG | GTAGTTTCGTGGATGCCACAG |
| *BECN1* | Human | TGCCGTTATACTGTTCTGGGG | CGTGTCTCGCCTTTCTCAAC |
| *CIDEA* | Human | TTATGGGATCACAGACTAAGCGA | TGCTCCTGTCATGGTTGGAGA |
| *CDKN2A (p16^Ink4a^)* | Human | GGGGTCGGGTAGAGGAGG | TCATCATGACCTGGATCGGC |
| *CDKN2A (p14^Arf^)* | Human | GGTTCTTGGTGACCCTCCG | TCAGTAGCATCAGCACGAGG |
| *PPARGC1A* | Human | TCTGAGTCTGTATGGAGTGACAT | CCAAGTCGTTCACATCTAGTTCA |
| *PRDM16* | Human | CTTCGGATGGGAGCAAATACTG | TCCACGCAGAACTTCTCACTG |
| *UCP1* | Human | AGAAGGGCGGATGAAACTCT | ATCCTGGACCGTGTCGTAG |
