## Supplementary figures and images for "*Cdkn2a* regulates beige fat maintenance through BECN1-mediated autophagy"

### Figure S1

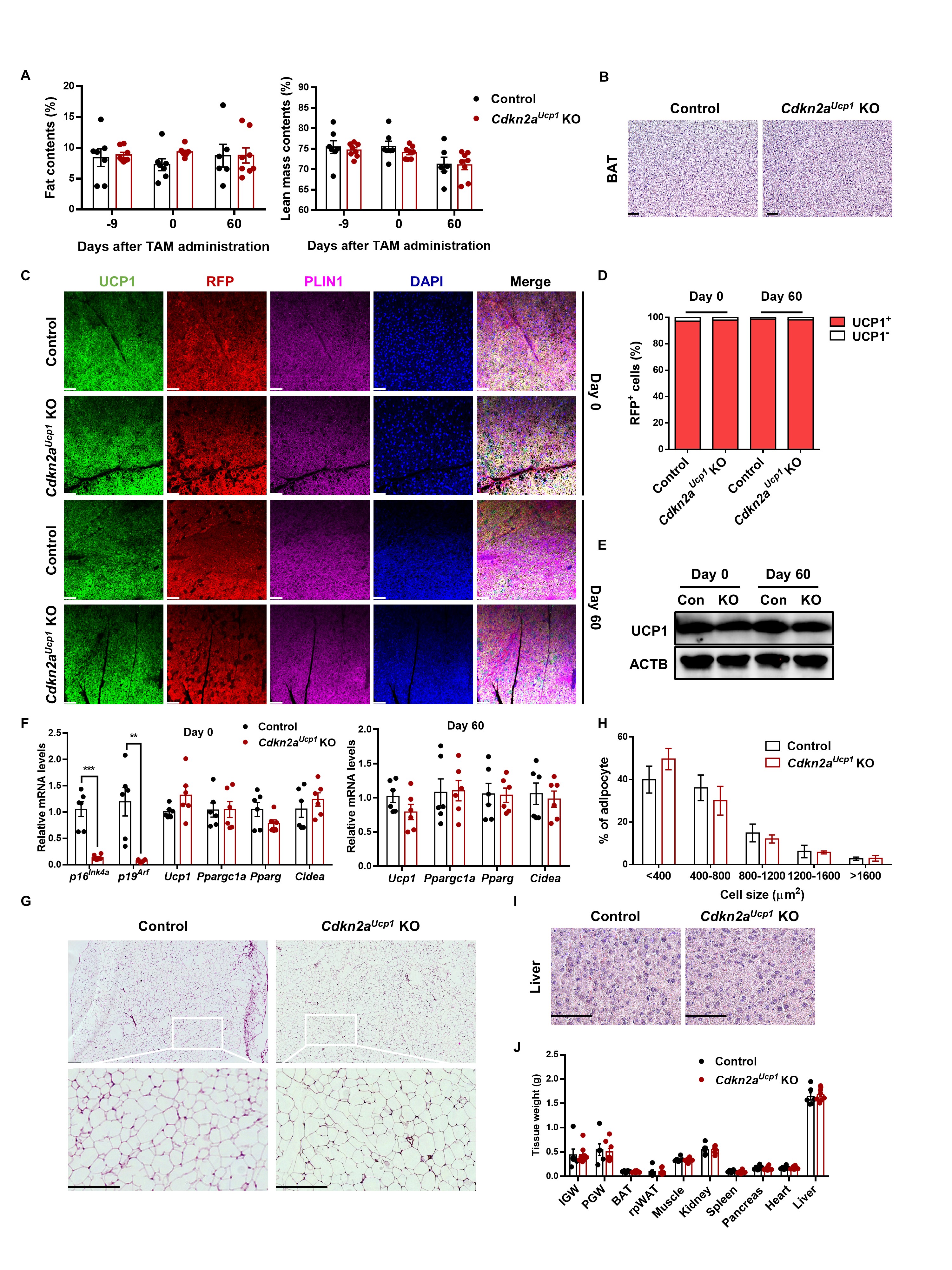

### Figure S2

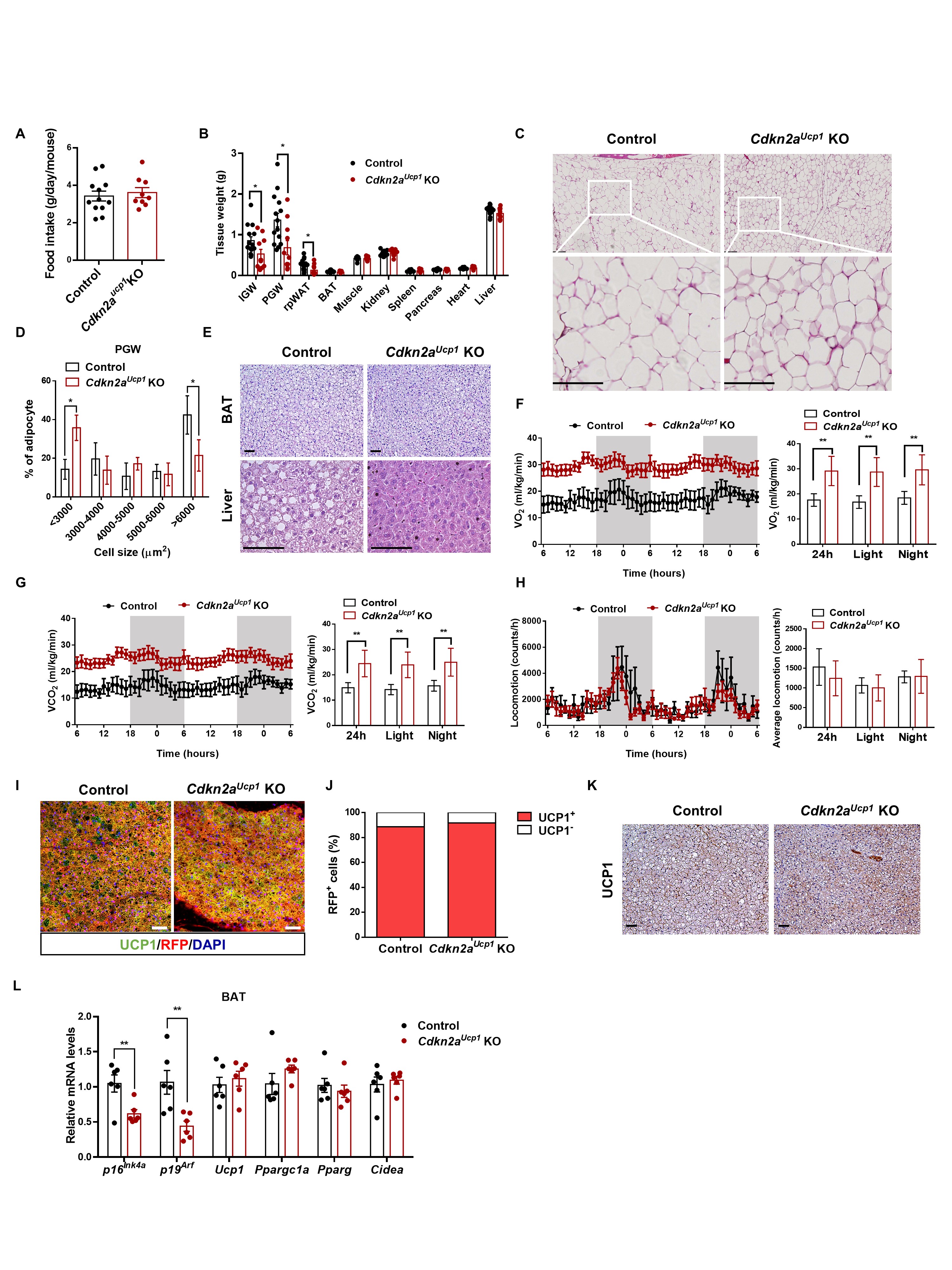

### Figure S3

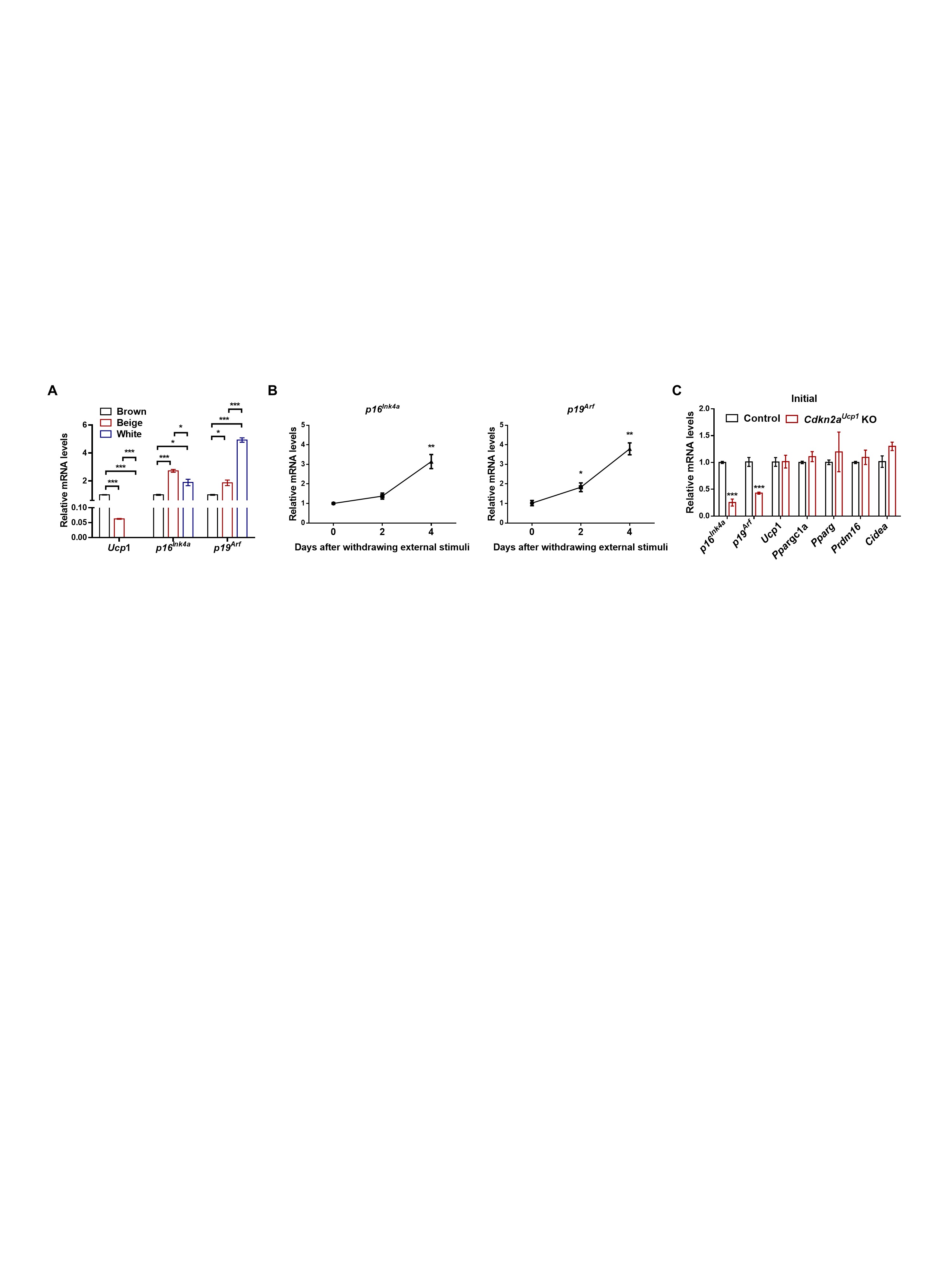

### Figure S4

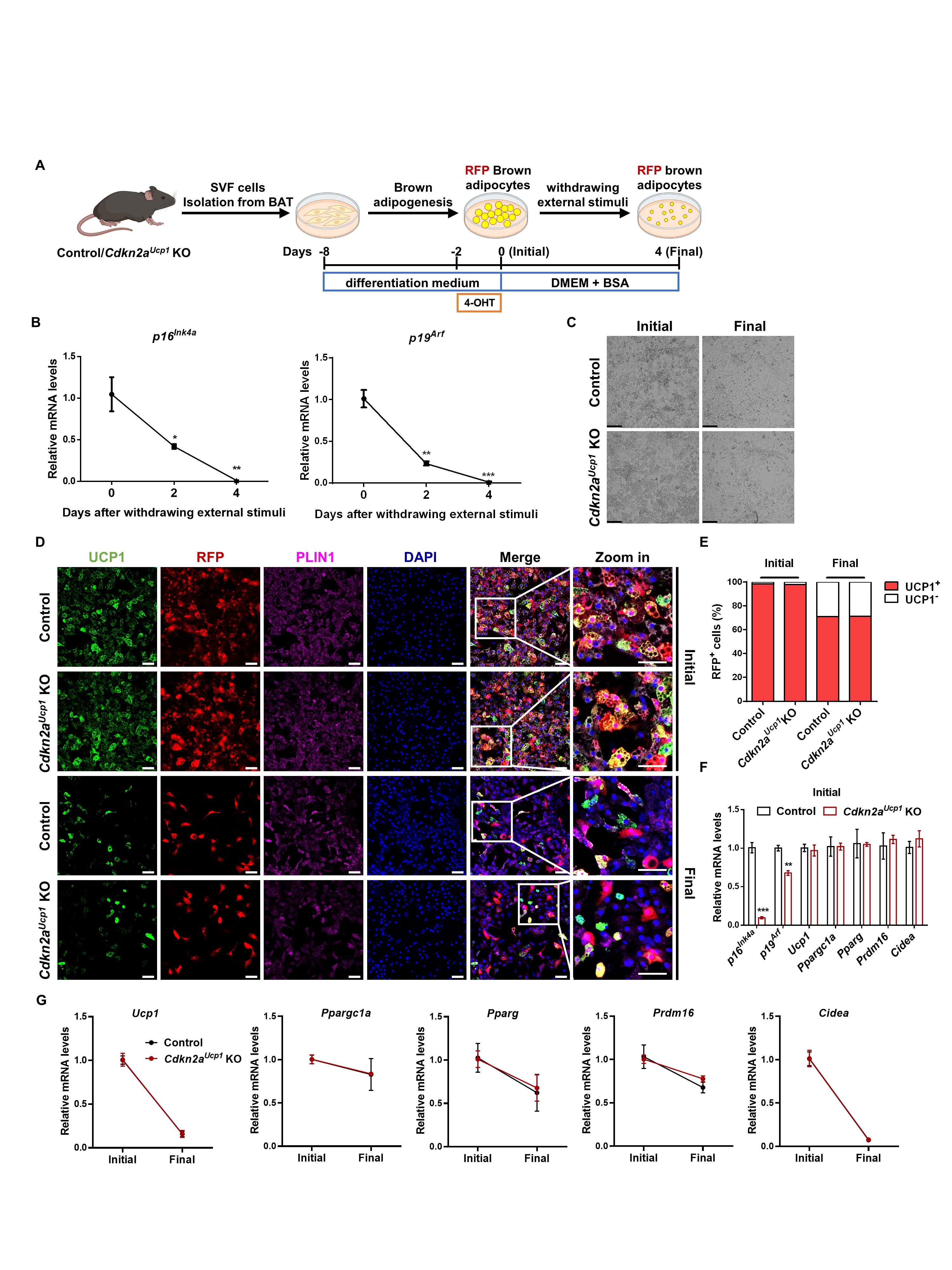

### Figure S5

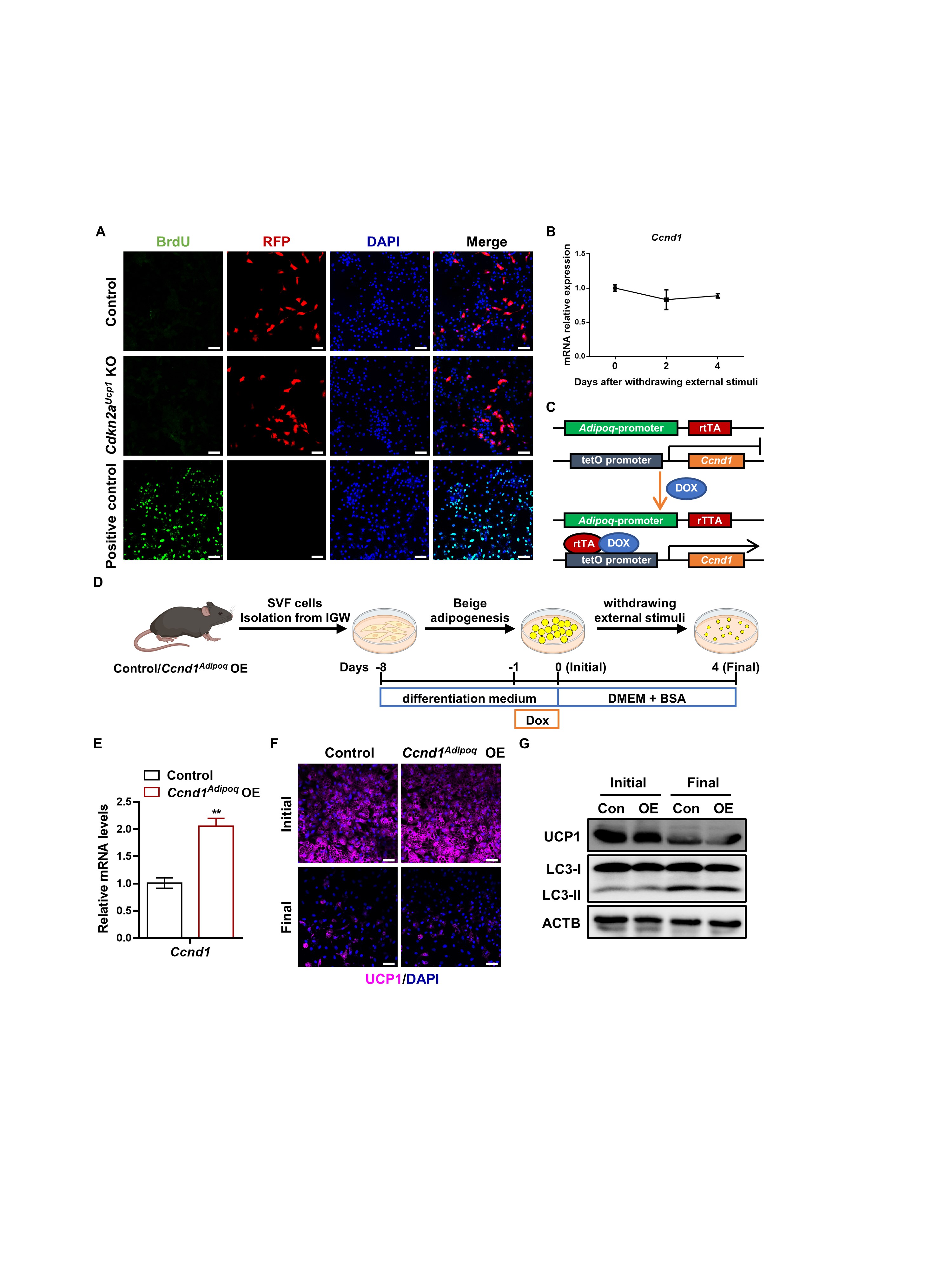

### Figure S6

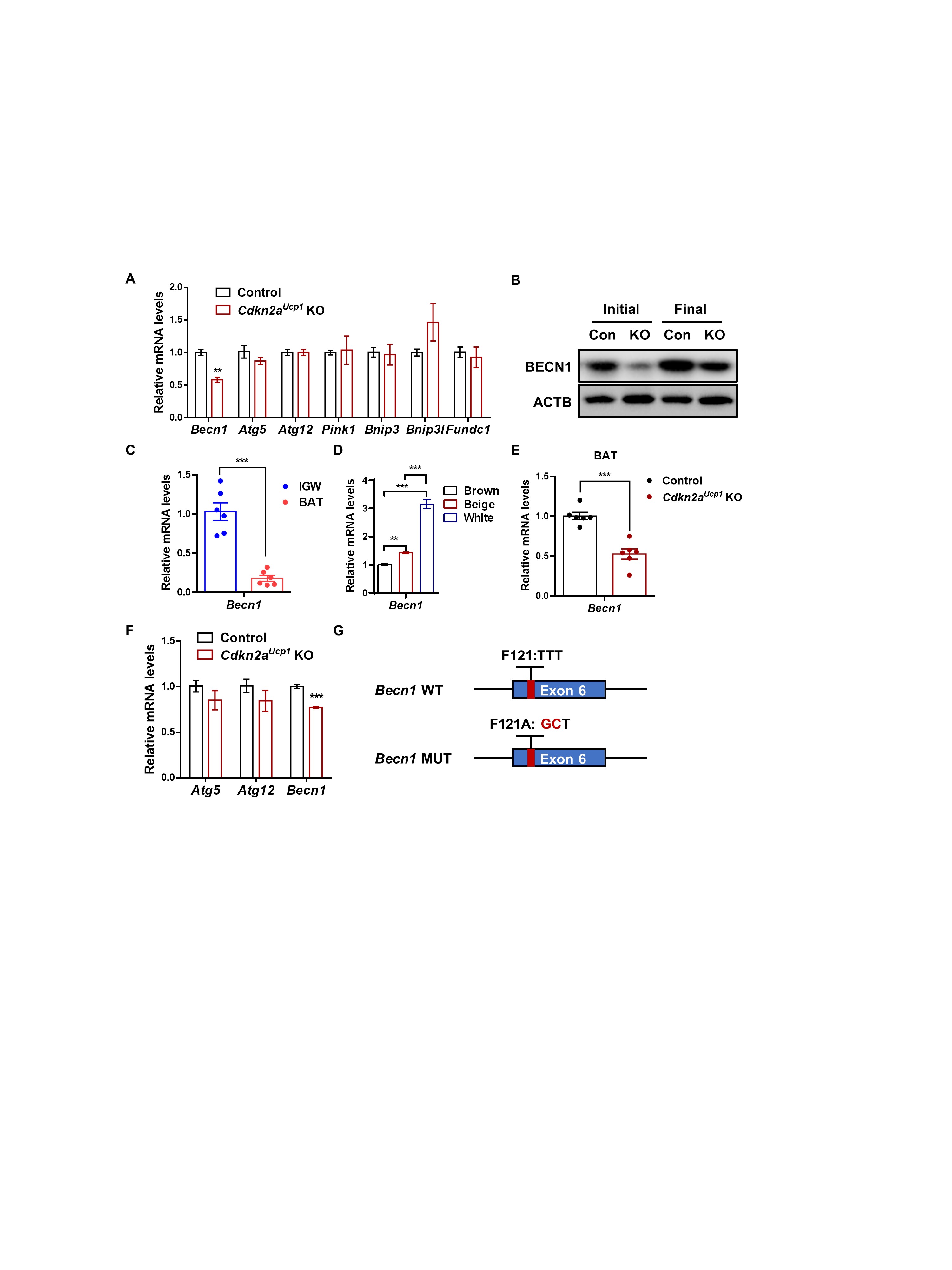

### Figure S7

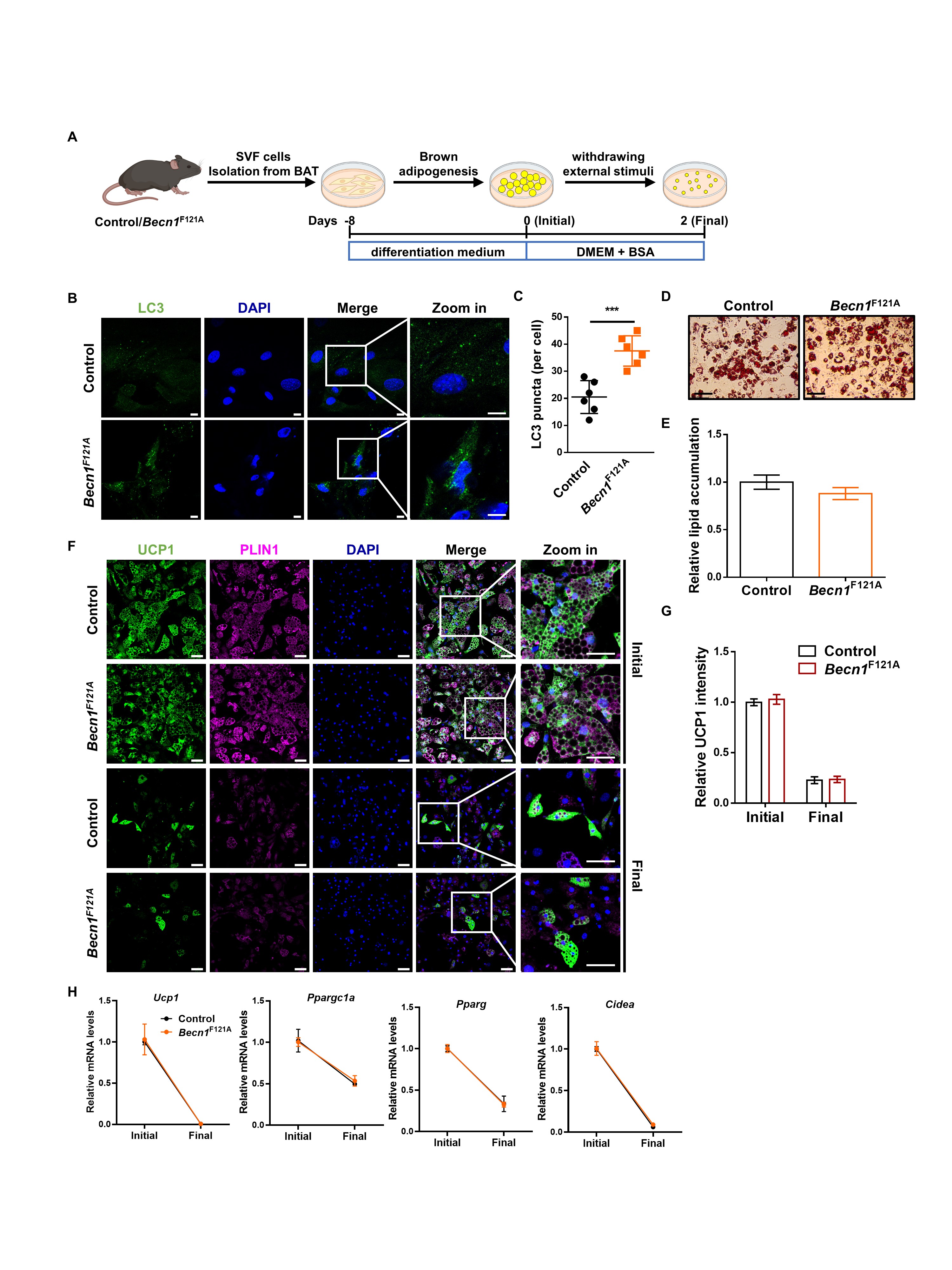

### Figure S8

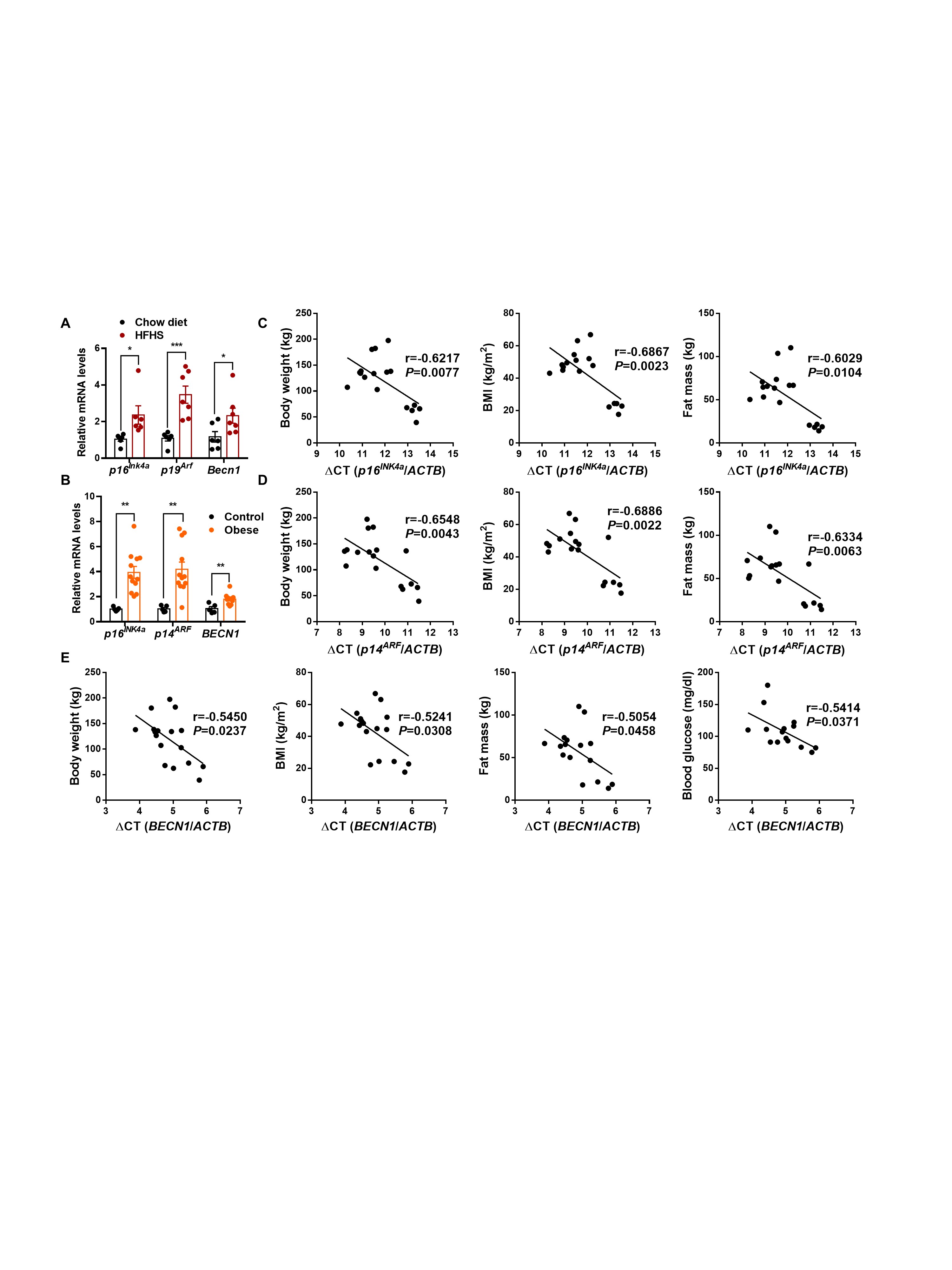
